# Dissecting targeted therapy resistance: Integrating models to quantify environment mediated drug resistance

**DOI:** 10.1101/156547

**Authors:** Noemi Picco, Erik Sahai, Philip K. Maini, Alexander R. A. Anderson

## Abstract

Drug resistance is the single most important driver of cancer treatment failure for modern targeted therapies. This resistance may be due to the presence of dormant or aggressive tumor cell phenotypes or to context-driven protection. Non-malignant cells and other factors, constituting the microenvironment in which the tumor grows (the stroma), are now thought to play a crucial role in both therapeutic response and resistance. Specifically, the dialogue between the tumor and stroma has been shown to modulate the response to molecularly targeted therapies, through proliferative and survival signaling. The goal of this work is to investigate interactions between a growing tumor and its surrounding stroma in facilitating the emergence of drug resistance. We use mathematical modeling as a theoretical framework to bridge between experimental models and scales, with the aim of separating the intrinsic and extrinsic components of resistance in BRAF mutated melanoma. The model describes tumor-stroma dynamics both with and without treatment. Calibration of our model, through the integration of experimental data, revealed significant variation across animal replicates in either the intensity of stromal promotion or intrinsic tissue carrying capacity. Furthermore our study highlights the need to account for this variation in the design of treatment strategies. ***Major Findings.*** Through the integration of a simple mathematical model with *in vitro* and *in vivo* experimental growth dynamics of melanoma cell lines (both with and without drug), we were able to dissect the relative contributions of intrinsic versus environmental resistance. Our study revealed significant heterogeneity *in vivo*, indicating that there is a diversity of either stromal promotion or tumor carrying capacity under targeted therapy. We believe this variation may be one possible explanation for the heterogeneity observed across patients and within individual patients with multiple metastases. Therefore, quantifying this variation both within *in vivo* model systems and in individual patients could have a significant impact on the design of future treatment strategies that target both the tumor and stroma. Further, we present guidelines for building more effective and longer lasting therapeutic strategies utilizing our experimentally calibrated model. These strategies explicitly consider the protective nature of the stroma and utilize inhibitors that modulate it.

**Precis:** Quantification of the environmental contribution to drug resistance reveals heterogeneity that significantly alters treatment dynamics that can be exploited for therapeutic gain.

**Financial Support:** Picco and Anderson: US National Cancer Institute grant U01CA151924.

Picco: UK Engineering and Physical Sciences Research Council (EPSRC grant number EP/G037280/1).

**Conflict of Interest Disclosure:** The authors declare no potential conflicts of interest.

## Quick Guide to Equations and Assumptions

The tumor is classified into two subpopulations, with respect to their sensitivity to the targeted inhibitor. *S* and *R* are, respectively, drug sensitive and drug tolerant populations. The stroma is divided into normal cells *F* (i.e. fibroblasts) and reactive cells *A* (i.e. cancer associated fibroblasts). The latter compartment represents fibroblasts in a transformed, secretory phenotype that promotes survival and tumor growth under drug treatment. We assume that *S* grows in the absence of treatment with growth rate *ρ*_S_, _R_grows under targeted treatment at rate ρ They share a carrying capacity *K*, representing the maximum packing capacity of the tissue where the tumor is growing. Targeted therapy (BRAFi) induces the stroma to switch to its reactive form at a rate *θ*. In turn reactive stroma (*A*) will promote tumor growth by an additional growth rate *η*. Upon removal of the targeted inhibitor, stromal renormalization occurs at rate φ and cancer cells are re-sensitised at rate *ξ*. The stromal-targeted inhibitor (FAKi) is assumed to reduce the stromal promotion by rate α These interactions occur dynamically in time (*t*) as defined by the following system of ordinary differential equations (ODEs):

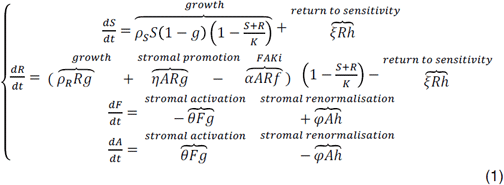

In addition we use the initial conditions:*S*(0) = *S* _0_,*R*(0)= *R* _0_,*F*(0) = *F* _0_, *A*(0)= *A* _0_. Note that,*g*(*t*),*h*(*t*) and*f*(*t*) are binary functions of time that allow for specific terms in the equations to be switched on and off, depending on treatment scheduling. Given a protocol calling for targeted therapy for the time interval 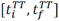 and FAKi for 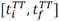 the binary functions are defined as follows

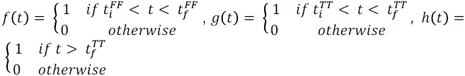

A useful measure of tumor burden control over a window of time [*t* _A_,*t* _B_] is the inverse of the area under the curve, defined as follows:

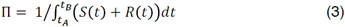

## Introduction

In the past decade many molecular targets of oncogenic drivers have been developed and approved for the treatment of pathway-specific cancers, in the hope that they could accompany or even replace highly toxic chemotherapeutic drugs [1-4]. Unfortunately this strategy turned out to be only partially successful, with strong initial responses often followed by relapse [3]. In an attempt to improve these poor long-term responses, combinations of multiple inhibitors (including immunotherapies) have been attempted [5-7]. Despite successes in concurrent inhibition of several pathways in preclinical models [8,9], it would seem that in the clinical setting, combination of targeted inhibitors does not offer cure, but can at best delay inevitable disease progression caused by the onset of drug resistance [2, 10, 11].

In an effort to understand why these treatment strategies fail, and how we might redesign better and more successful treatments, we must embrace the reality that cancer is a complex evolving system. Because cancer is an evolutionary disease, it can evolve strategies to override or circumvent the action of a given inhibitor. These strategies include producing secondary mutations [11] or exploiting pre-existing genetic heterogeneity. However, mutations alone are not sufficient to explain the often rapid timescale over which cancer stops responding to therapy [12]. Recent evidence suggests that cancer is able to co-opt the surrounding stroma to create an environment that can facilitate treatment escape [13, 14]. This phenomenon is termed Environment-Mediated Drug Resistance (EMDR) [12], and includes several processes ranging from cell-adhesion mediated drug resistance [15-17] to therapy-induced secretomes such as IGF, HGF, TGF-beta [8, 18] and fibronectin [19]. The mechanisms of context-driven resistance we consider here are shared across a variety of solid tumors characterized by aberrations in growth-control signaling and a high level of interaction with the surrounding tumor microenvironment. Our primary focus here is on BRAF mutated melanoma. A particular instance of EMDR in melanoma is represented by the action of cancer associated fibroblasts (CAFs) that create a habitat favorable for drug tolerance and tumor growth. The environmental remodeling includes deposition of extracellular matrix (ECM) components, upregulation of growth factor production, intensification of paracrine signaling between the stroma and the tumor cells, and rewiring of the tumor cells’ proliferative and survival signaling via integrin binding [12]. The effect of this transformed habitat on the cancer and stromal cells is transiently induced by application of the targeted drug and is mostly reversible [20]. Given the transient nature of EMDR there may be an opportunity to modulate it through treatment holidays by allowing renormalization of the stroma to occur – potentially facilitating a better overall treatment outcome. Additionally, preliminary investigations have shown benefits in inhibiting stromal-derived processes, such as elevated FAK signaling [13]. Dual targeting of tumor and stromal processes represents a promising strategy for better management of BRAF mutated melanoma.

Understanding this complex interplay between tumor and host cells undergoing treatment is ideally suited for mathematical and computational models. Recently, several theoretical studies have addressed the role of the environment in facilitating drug resistance. Mumenthaler et al. have studied how gradients of nutrients and drug concentration modulate the fitness of drug-sensitive and drug-resistant cell lines, and eventually determine recurrence [21]. Sun et al. modeled the environmental adaptation to drug treatment via drug-induced resistance factors that modulate the growth dynamics of metastatic disease [22]. Silva et al. and, more recently, Robertson-Tessi et al. modeled microenvironmental heterogeneity, specifically the regulation of metabolism, to understand the evolutionary dynamics driving treatment response and leading to resistance [23, 24]. A significant literature already exists for mathematical models of intrinsic resistance in cancer progression and response to treatment. Lavi et al. offer a comprehensive review of models of cancer resistance [25]. However, the focus of the majority of these models is limited to intrinsic chemotherapy resistance [26]. Models that integrate the role of the stroma, which is key in the emergence of resistance to targeted therapeutics, are less well developed, but are beginning to emerge. Many studies analyze the dynamics emerging from tumor-immune interactions [27-30]. Fewer mathematical models specifically describe interactions between cancer and stromal fibroblasts and their role in drug resistance [31-34]. To our knowledge, the problem of separating intrinsic resistance from EMDR, through the dynamics of response to targeted therapy, has not yet been addressed.

Here we present a first minimal model of tumor-stromal interactions, which aims to bridge the growth dynamics of cancer from *in vitro* and *in vivo* experimental models. Specifically, by using exponential growth dynamics from cells growing *in vitro* we can calibrate baseline unconstrained growth dynamics. Then using the same cancer cell line *in vivo* (mouse allograft) we can capture the saturation dynamics. Using these two experimental model systems, under treatment, we can then quantify the relative contribution of the environment to tumor growth.

Our calibrated model describes the baseline growth dynamics and the relevant tumor-stromal interactions determining growth and response to treatment. This, in turn, allows a fuller exploration of the role of stroma in the promotion of drug resistance, which we propose is critical for the design of optimal treatment strategies. To this end we will explore treatment schedules that exploit tumor-stromal interactions to limit and/or delay the emergence of EMDR. Our study gives preliminary guidelines for building more effective and longer lasting therapeutic strategies, including dose fractionation and timing.

### Materials and Methods

A common paradigm for the treatment of advanced stage BRAF-mutated melanoma includes targeted therapy in the form of a BRAF inhibitor (BRAFi), such as vemurafenib, recently approved for patients carrying the V600E mutation [35]. Kinase inhibitors such as vemurafenib specifically block a molecular pathway that the cancer cells are strongly dependent on, resulting in reduced toxicity for the whole body and increased specificity for the tumor. While this treatment can keep the cancer in check for many months, the disease will eventually recur. Having identified the environment as a key factor in therapy failure [12], alternative blockades of stromal-derived processes are actively being investigated. Here we specifically model FAK inhibition (FAKi) that has proved effective in the pre-clinical setting [13].

We propose a model of EMDR for molecularly targeted cancers. Figure 1 shows a schematic of the interactions between key players in our system: cancer cells classified as either sensitive or tolerant to the targeted drug (*S* and *R*, respectively), and stroma cells in normal or reactive form (*F* and *A*, respectively). The *R* compartment accounts for an initial intrinsic resistant cancer population as well as cells that are transiently drug-tolerant through the action of EMDR. It is worth noting that this “catch all” compartment does not correspond to a single biological phenotype or genotype, however, it allows us to analyze growth regimes with and without targeted treatment, and most importantly to quantify the relative contribution of the environment to tumor growth dynamics under treatment. Significant bacterial literature indicates the existence of persister phenotypes which are tolerant to a number of antibiotic agents and yet do not appear to be driven by genetic changes [36]. Very recently, such populations of ‘cancer persister cells’ have been discovered in an EGFR+ lung cancer cell line [37, 38]. However, in the absence of more detailed data, we develop a simplified model with an initial *R* population that includes cells derived from any of these mechanisms, and allow all cells to return to sensitivity, irrespective of resistance mechanism.

**Figure 1.**
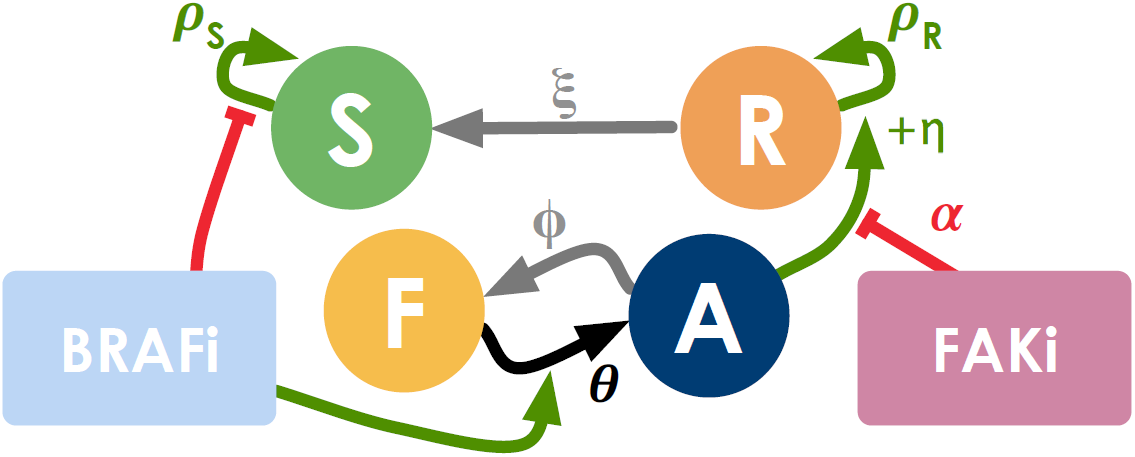
Interactions hypothesized in the compartmental model: positive interactions are represented with green arrows, negative ones with red flat ends. The BRAFi (targeted to the tumor) inhibits growth in the drug-sensitive portion of the tumor (S) and induces activation of normal stroma (F). In turn, reactive stroma (A) promotes growth in the drug-tolerant portion of the tumor (R). The stroma-targeted inhibitor FAKi dampens the effect of stromal-induced growth promotion. Upon removal of BRAFi the tumor reacquires sensitivity to the drug and the stroma renormalizes (grey arrows).

The interactions between the cell compartments, modulated by the two drugs (BRAFi and FAKi), are defined by a set of ordinary differential equations (ODEs) (1) discussed in more detail in the guide to equations. A key advantage of this simple model is that it can incorporate data from both *in vitro* and *in vivo* experimental models.

Figure 2 shows the experimental data for BRAF mutated melanoma cell lines 5555 and 4434. These cells were both cultured *in vitro* (Figure 2A) and injected *in vivo* (Figure 2B). Growth was observed over time, both in the absence of drug and under treatment with PLX4720, a BRAF inhibitor. We can adapt the model (1) to represent each one of these experimental conditions. Table 1 shows a summary of the experimental conditions and corresponding models. Starting from the *in vitro* experimental setup, corresponding to a simplified system of equations with fewer unknown parameters, we obtain parameter estimates by data fitting and consequently use these values for the data fitting of the *in vivo* experimental setup. In doing so we significantly reduce the number of unknown parameters for each fit, as well as the risk of overfitting.

**Figure 2.**
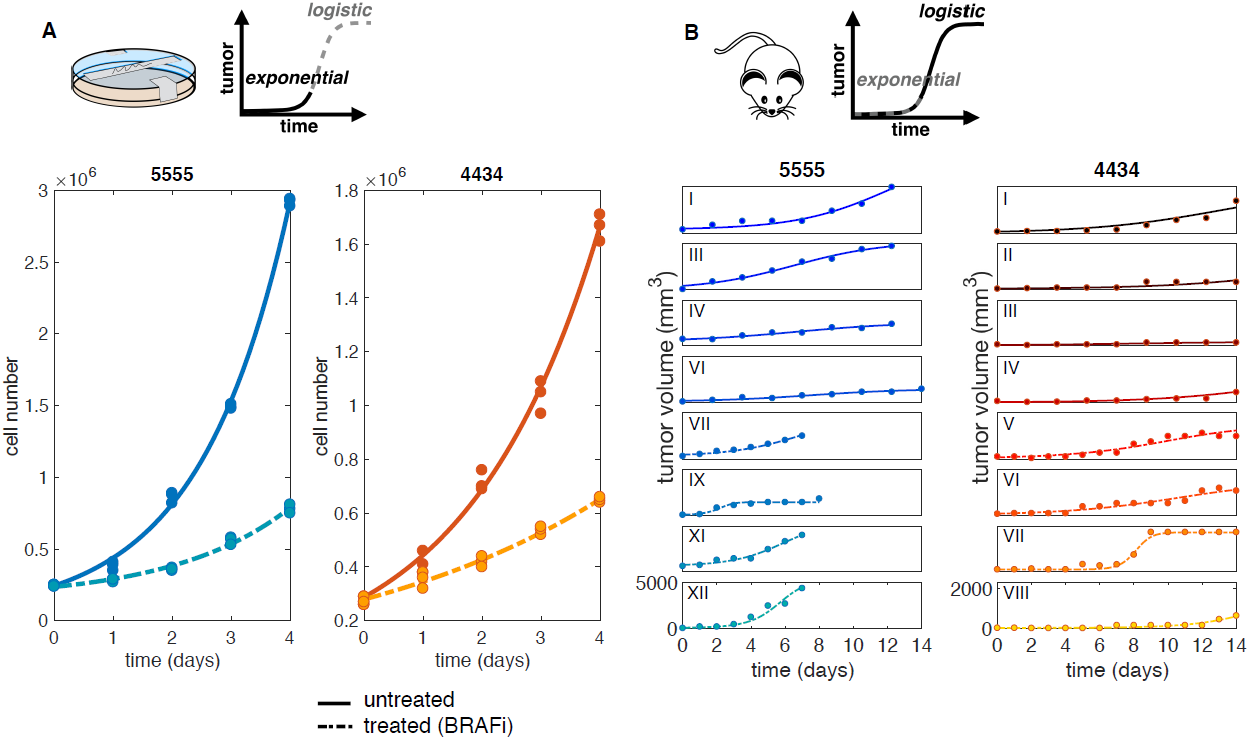
Unpicking the relative contributions of intrinsic resistance and extrinsic environment conferred tolerance (EMDR). **A.** *In vit*r*o* data and fit. For each condition we obtained one estimate that best fits the three replicates at the same time. **B.** *In vivo* data and fit. Data consist of six and four untreated 5555 and 4434 mice, respectively, and six and five BRAFi treated 5555 and 4434 mice, respectively. Only few representative mice are shown. For each condition the model is fitted individually to each replicate (mouse). Solid and dashed lines correspond to untreated and BRAFi treated tumor, respectively. Note different y axis scale for the two cell lines. Data from [13].

The *in vitro* setup (with a time scale on the order of a few days, Figure 2A) can be represented by an exponential growth regime, and lacks the stromal component. This corresponds to reducing system (1) for small time *t* with *F* _0_= 0, obtaining:

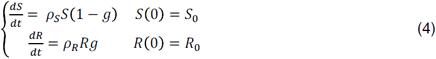

where the only unknown parameters are: R_0_ S_0_ ρS for the untreated case (*g* = 0), and R_0_, S_0_, ρ _R_ for the treated case (g= 1). Parameter estimation for these triplets is carried out by Approximate Bayesian Computation, which builds a discrete approximation of the posterior distribution. Data are fitted to the analytical solution of (4). Analytical solutions are reported in Table 1 and a detailed description of the estimation method is reported in the supplementary material. Figure 3A shows the marginal distributions for the growth rates of each cell line. Comparing the estimates for control and treated conditions, we see a reduction in growth rate for the treated cancer. The deficit in growth rate reveals that under drug treatment the *R* population, irrespective of the mechanism of resistance, exhibits slower growth compared to the *S* population in untreated conditions, consistent with the previous literature [20].

**Figure 3.**
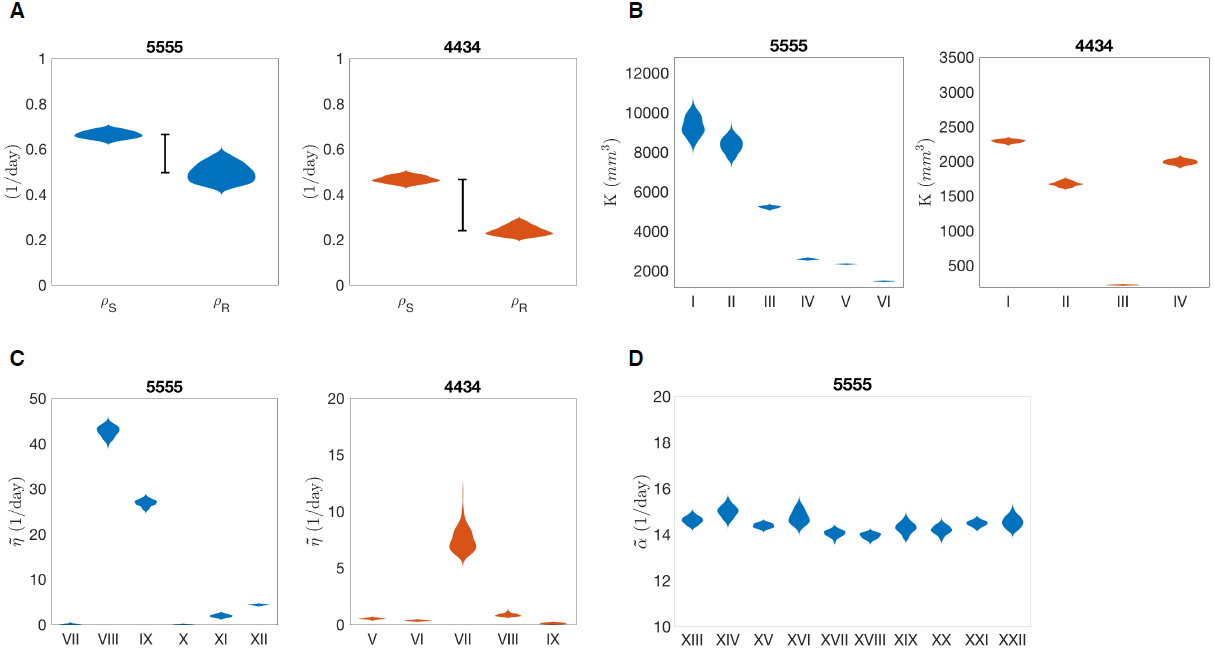
Approximated posterior distribution of estimated parameters. **A.** Estimates for cancer growth rates: ρ _S_ (untreated cancer), ρ _R_’ (treated cancer). The black bar highlights the fitness cost of intrinsic resistance. **B.** Estimates for *K* reveal heterogeneity of carrying capacity across mice. ρ _S_ from previous estimate (Table 2). **C.** Estimates for 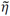 reveal heterogeneity of stromal-derived protection found *in vivo*. θ= 0.03 1/*day*. ρ _R_ from previous estimates (Table 2). **D.** Estimates for 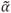 for ten 5555 mice treated with BRAFi and FAKi combination. θ = 0.03 1/*day*.,ρ _R_= 0.49539 1/*day*. 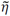 =12.67 1/*day*(average of previous estimates, Table 2).

We assume that the growth dynamics of the cancer cells treated *in vitro* can be solely attributed to pre-existing drug tolerant subpopulations. On the other hand, in order to quantify the role of the environment on the dynamics of resistance, we turn to the mouse allografts. When the same cell lines are injected in mice, growth is significantly constrained and experiments cover a longer time scale (Figure 2B). The observed dynamics are more accurately captured with a logistic growth regime, as described by:

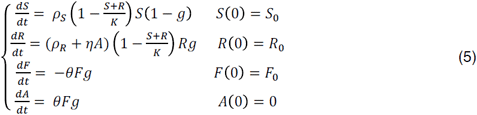

By assuming that cells from the same cell line grow at the same exponential rate in an unconstrained environment, we are able to use the growth rates estimated from the *in vitro* data (i.e. and ρ _S_ and ρ _R_) to help calibrate the parameter estimates for the *in vivo* model.

By fitting the model to the untreated mice data we obtain estimates for the parameters R_0_,S _0_,*K*.Note that in the absence of treatment (g= 0), the equations for the tumor and stromal populations are decoupled, therefore the estimate of tissue carrying capacity (K) is independent of the quantification of interacting stromal cells. However, *K* is intrinsically dependent on nutrient constraints as well as the packing capacity of the tissue. Indeed, variations of this quantity are captured in the range of estimated values (see Figure 3B and Table 2). Posterior distributions are wider in mice with higher values of carrying capacity (e.g. mice I, II cell line 5555). For low *K*, logistic curves reach carrying capacity within the time window of the *in vivo* experiments. Curves with higher *K*, however, have a later inflection point and their characteristic shape is not captured in the same time window, resulting in more uncertain estimates. It is worth noting that for some mice, the data do not capture the saturating dynamics, as the experiment had to be interrupted due to animal welfare (for details on the original experiments see [13]).

For the treated mice setup (g= 1), the equations are coupled. We can solve the last two equations of (5) analytically, to write *A* as a function of F_R_ and θ. Defining 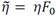, we reduce the parameter number in the analytical solution of (5). At this stage, experimental quantification of the rate of stromal activation is not available; the before estimates for the parameters R _0_,S _0_,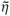 will be carried out with a range of θvalues. We observed high sensitivity of the estimates of 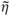 to variations in this experimentally undefined parameter θ (Figure S3). Figure 3C shows the estimated values for 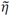 for each mouse. This reveals considerable variation in the stromal support across mice, hinting at an underlying heterogeneity in stromal habitats and activation. Since estimates of *K* and ñare dependent on the previously estimated growth rates (ρ _S_ and ρ _R_ respectively), the ABC estimation was run for values of growth rates within the range captured by the fit to the *in vit*r*o* data (see Table 2). The resulting posterior distributions varying in relation to the growth rates are shown in Figure S1 and S2, respectively. The variation in response to BRAF inhibition across replicates could be the result of underlying heterogeneity either in tissue carrying capacity or in stromal support, or both. Our estimation protocol for the stromal promotion parameter 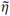 makes use of an average carrying capacity *K* previously estimated. However the variation across replicates could also be explained by variation in carrying capacity. Therefore we further investigated the BRAFi treated mice data, to infer the posterior distribution of 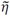 as Kis varied and *vice ve*r*sa*. Figure S4 shows the resulting posterior distributions in the (k, 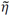) space for a sample mouse. The posterior distribution of *K* is highly sensitive to the variation of 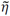 (see yellow violin plots), and *vice ve*r*sa*. However the best overall fits of 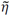 and *K* are located in the same region of the space. This means that for a given mouse we can unequivocally identify a combination of values for the carrying capacity and stromal support that best explains the data.

Finally, we can quantify the inhibiting action of the stromal-targeted drug in the form of 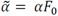 fitting data from mice treated with both BRAFi (PLX4720) and FAKi (PF562271) to the following version of the model:

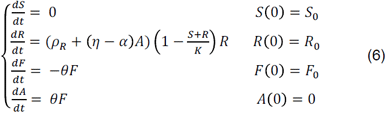

Despite the variability of responses across replicates (see data and fits in Figure S5), the resulting estimates for 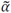 show little variation (Figure 3D and Table 2). This implies that the variability in treatment response may be attributed to the heterogeneous stromal composition of the tissue (highlighted in Figure 3C), as opposed to the efficacy of the stromal inhibition.

### Results

Calibrating our model across *in vit*r*o* and *in vivo* data allows us to gain insight into the dynamics of the system that a qualitative analysis of these experiments cannot capture. Figure 3A shows the marginal posterior distribution for growth rates *ρ*_S_ and *ρ*_R_, with a reduction of the latter quantifying the impact that drug tolerance has on proliferative capacity.

Comparing *in vit*r*o* and *in vivo* dynamics allows us to assess the relative contribution of the environment to drug resistance. This analysis revealed significant heterogeneity across replicates (mice), both in terms of tissue carrying capacity, and stromal protection (Figures 3B and 3C). This heterogeneity translates to a high variability of response to treatments that target both the tumor and the stroma, despite the apparent more homogeneous inhibitory effects of the stromal-targeted drug (Figures S5 and 3D).

Analysis of the ODE model with the combination treatment of BRAFi and FAKi (equation (6)) gives insight into the dynamics of the system as a function of stromal promotion and tumor growth rate. Specifically, we can discriminate two distinct cases:

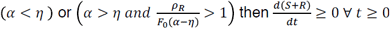

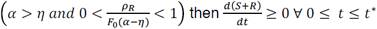

where 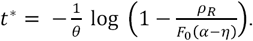

In the first case, either the stormal promotion is too strong to be compensated by the FAKi, or the stromal promotion is weak, but the tumor growth rate is elevated. Then the overall tumor burden is monotonically increasing, although bounded by the carrying capacity, and therapy is ineffective. In the second case, when stromal promotion is weak and the tumor growth rate is reduced, then the therapy is effective provided that it is administered for a sufficiently large period of time.

As an example, consider the cohort of 5555 BRAFi-treated mice (VII through XII) and using the parameterized model (1), with α taken as the average of the previous estimates (see Table 2), we can sub-classify the mice. According to our estimates, mice VII, X, XI, XII fall into case 2, meaning that with a combination of BRAFi and FAKi it is possible to achieve control as long as we treat past time *t*^*∗*^. On the other hand, mice VIII and IX fall into case 1, hence the tumor is always growing under treatment, eventually reaching carrying capacity. Figure S6 and S7 show a simulated treatment combination of BRAFi + FAKi calibrated on two representative mice, case 1 and case 2, respectively.

For a tumor-stroma system falling into case 1, recurrence is inevitable, but may be delayed with alternative scheduling strategies. Given that the phenotypic changes underlying EMDR are transient and reversible upon drug removal, we hypothesize that the introduction of drug holidays could significantly improve treatment response and recurrence times. Intermittent application of vemurafenib has proved to be successful in melanoma xenograft models [20] and ongoing clinical trials are testing intermittent versus continuous dosing of a combination of BRAF and MEK inhibitors (NCT02196181). However, we believe that a mechanistic and quantitative approach to treatment scheduling can improve the success of the otherwise empirical approach that these studies offer. We therefore systematically explored the space of holiday versus treatment days of an intermittent schedule treatment with BRAFi, combined with continuous FAKi.

Specifically, the targeted inhibitor is administered during the time windows 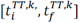 for *k* ϵℕ ^R^.with 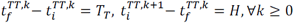. That is, we consider treatments of fixed duration T_U_, with the time between the end of one treatment and the start of the next treatment being fixed at H. Figure 4 shows the treatment outcome in the holiday vs. treatment space (H, T_U_), where the outcome of each treatment strategy over the time frame of 0,70 *days* is quantified with ∏, defined in (3). This reveals that the region corresponding to tumor burden minimization (Π maximisation) is concentrated around the line *H* = 2T_U_. Intuitively this means that the length of holiday needed to renormalize the system is proportional to the pulse of treatment. Additionally, it indicates that longer treatment holidays are more effective at controlling tumor burden, while the total number of treatment days is reduced.

**Figure 4.**
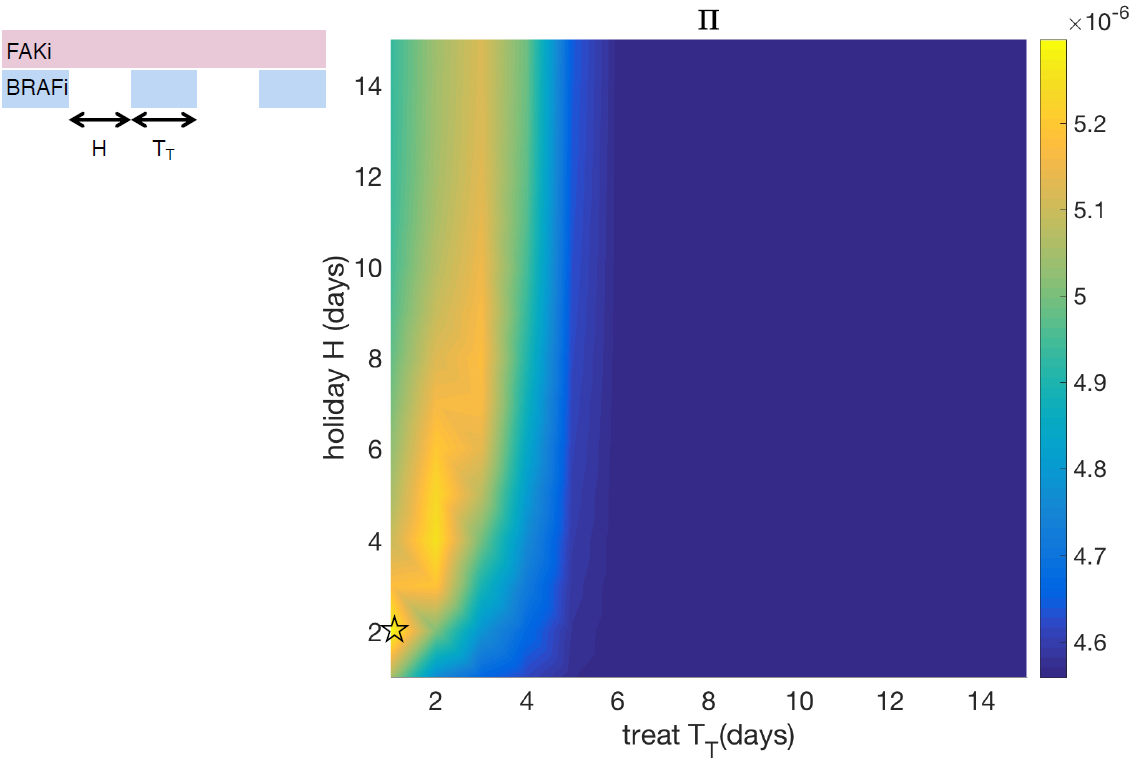
Exploration of treat/holiday space for intermittent BRAFi combined with continuous FAKi to maximize control of tumor burden. Surface plot of Π(see equation (3)) in the treat/holiday space. Model parameterized on mouse IX of cell line 5555.,*ρ*_S_ = 0.66325 1 *day*,*ρ*_R_’ = 0.49543 1/*day*, *K* = 4818.62 *mm*^3^, 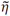 = 26.876 1/*day*, 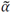 = 14.4 1/*day* θ= 0.03 1/*day*, ξ = 0.01 1/*day*, φ=11/*day*, *S* _0_ = 48 *mm* ^3^, *R* _0_= 12 *mm*^3^, *F* _0_= 60 *mm*^3^, *A* _0_= 0 *mm*^3^. The star indicates the treatment schedule simulated in Figure 5.

Figure 5 shows the temporal dynamics for one of the best combination treatment schedules predicted by our model. FAKi is continuously administered, and helps control the tumor burden when EMDR sets in, whereas BRAFi is periodically for 1 day, then off for 2 days. This treatment induces only minimal stromal activation and delays progression by approximately ten days when compared to the untreated tumor. When compared to the continuous treatment, this intermittent treatment delays progression by approximately twenty days, while using a third of the amount of BRAFi. Although this study does not explicitly account for drug toxicity, total dose reduction is a desirable outcome, especially in the case of combination therapy, where resulting toxicity might be a significant issue.

**Figure 5.**
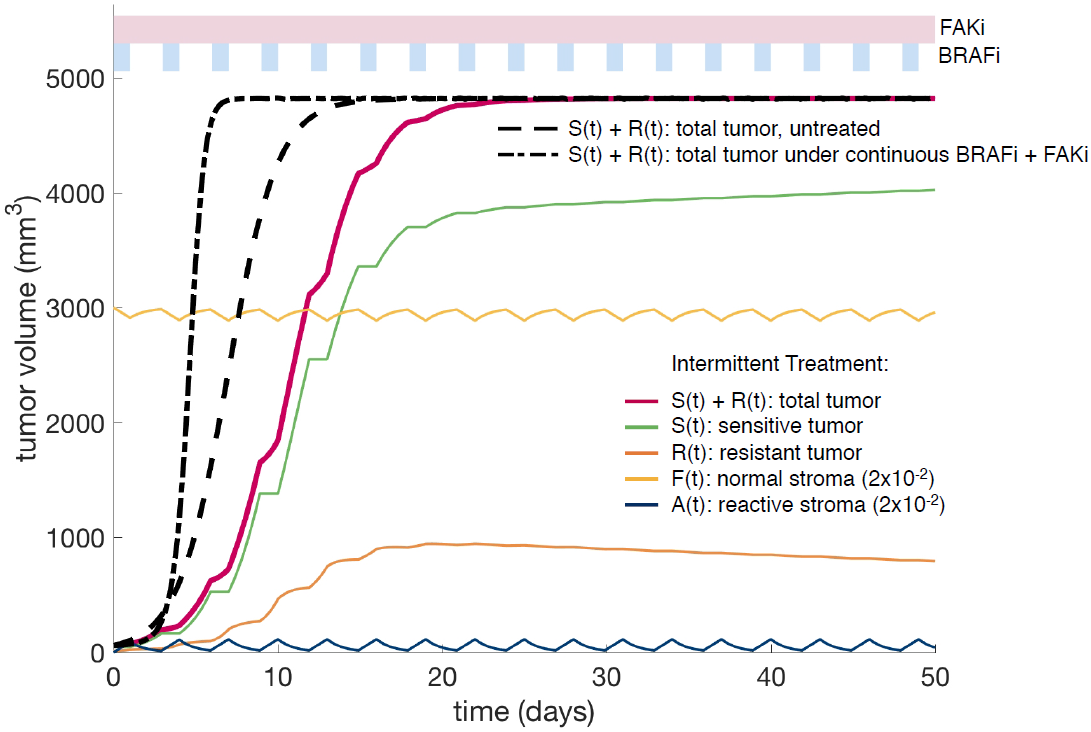
Example of combination therapy schedule (BRAFi+FAKi) to exploit tumor-stroma interactions. The model is parameterized as reported in Figure 4. Blue and pink bands above the graph indicate the BRAFi and FAKi administration windows. The BRAFi is intermittently administered for 1 days with 2 days holiday. The FAKi is continuously administered. This treatment schedule delays the disease progression by approximately ten days (compare red and dashed black lines) while using a third of the BRAFi dose. The green and orange lines show the breakdown of the total tumor burden in sensitive and resistant population, respectively.

## Discussion

Molecularly targeted therapies for cancers with known driver mutations are extremely effective for six to eight months (e.g. vemurafenib for BRAF V600E melanoma [39]) and are accompanied by lower toxicity when compared to cytotoxic chemotherapeutic agents [3]. However, with continuous and prolonged treatment, the emergence of drug resistance seems to be inevitable. Upon removal of the targeted drug, due to relapse, a typical disease flare is observed (e.g. EGFR mutated lung cancer treated with a combination of tyrosine kinase inhibitors [40]), suggesting that the treatment has somehow selected for a more aggressive clonal population in the tumor. However, subsequent treatment with the same inhibitor often leads to an additional response [41, 42], suggesting that selection of resistant clones alone cannot explain this disease etiology. The environment is now considered an important source of non-intrinsic drug resistance mechanisms [43], collectively referred to as Environment Mediated Drug Resistance. Since the changes accompanying EMDR are considered transient and therefore reversible, the possibility of regulating EMDR dynamics with smarter treatment scheduling is promising. However, a necessary first step towards the design of such treatment strategies is a more quantitative understanding of the interactions and dynamics occurring between the tumor and the stroma.

*In vit*r*o* model systems can accurately quantify temporal tumor growth and treatment response in controlled environments, whereas *in vivo* models more readily capture the native environment that is directly relevant to patients. However, both of these are models of human disease and only capture specific aspects of reality over very specific spatial and temporal scales. The ODE model we develop here bridges between these experimental scales, in order to integrate relevant information from each of them.

Starting from analysis of BRAF-mutated melanoma cell lines, we quantified the baseline dynamics of cancer cells in a uniform nutrient-rich environment. By comparing the growth rates of cells untreated and treated with the BRAF inhibitor, we were able to quantify the overall reduction of growth under drug application. Our model facilitates this analysis by classifying the cancer into two separate populations, growing with or without drug (*R* and *S*). Then using Approximate Bayesian Computation we calculate plausible regions of parameter values. This type of estimation can be particularly useful when assessing the error in fitting. Our parameter estimation method does not make assumptions on the initial conditions, and *R*_0_ and *S*_0_ are included in the parameters to be estimated. Consequently, the model is agnostic to the mechanisms producing the initial resistant population, *R*_0_. These mechanisms could be EMDR-related as well as epigenetic or non-autonomous. However, at this stage, no data are available to distinguish between these instances of resistance, and we group all cells that grow under drug treatment in the *R* compartment, irrespective of the underlying mechanisms of resistance.

Subsequent analysis of data from mice xenografts implanted with the same cell lines allowed us to identify the relative contribution of the environment to drug resistance. This analysis revealed heterogeneity in both the local tissue carrying capacity and in the stromal promotion of tumor growth. This heterogeneity maybe one possible explanation for the spectrum of response observed across patients. In the context of metastatic disease, with tumors seeded across a variety of tissues, heterogeneity in stromal composition could be an important discriminating factor in the success of a systemic treatment. Therefore, quantifying this variation in individual patients could have a significant impact on the design of future treatment strategies that target both the tumor and stroma.Within the current experimental and modeling framework, assessing the strength of stromal protection is non-trivial. This quantity is dependent on the abundance of the interacting stroma (we could only estimate the overall promotion rate (ñ=ηF _0_) as well as the speed of drug-induced stromal activation (we found high sensitivity to parameter θ). At the same time, with the available data, we can explain the variability of responses across mice by variation in carrying capacity and/or stromal promotion (Figure S4). Further investigation of the heterogeneity that our study revealed would require additional experimental quantification of these stromal-related processes. This would, in turn, allow us to address the main shortcoming of the current model, namely the high sensitivity of the estimate of stromal protection to the parameter θ (Figure S3).

Analysis of our ODE model revealed that the degree of stromal protection η, and cancer proliferation *ρ*_R_ drug treatment, are key in discriminating between responses to the combined action of inhibitors targeting tumor and stromal processes (BRAFi and FAKi, respectively). We found that for slower growing tumors it is possible to keep growth in check provided treatment with BRAFi is applied for a sufficient period of time. Conversely, for fast growing tumors or elevated stromal protection, the tumor burden increases, despite the administration of the inhibitors. However, for these tumors we can exploit the transient nature of the EMDR-associated mechanisms and delay progression. Specifically, scheduling treatment holidays for the mouse-(patient-) specific calibrated model would allow for renormalization of the system directly translating into better disease burden control. We used our parameterized ODE model to explore the space of intermittent treatment strategies, with the hope of improving response in cancers falling into the treatment refractory category. Neglecting toxicity of targeted drugs, we searched the space of holiday versus treatment length for intermittent BRAFi application, combined with continuous FAKi. We found that most effective tumor control is achieved with short BRAFi treatment pulses and longer holidays, requiring significantly less inhibitor, when compared to the continuous treatment.

It is worth noting that in optimizing the treatment schedule for these inhibitors, we are only modulating the dynamics by reducing the emergence of EMDR. This allows us to delay recurrence by approximately ten days. If we were to combine this strategy with a cytotoxic treatment, such as chemotherapy, which provides additional reduction of the tumor burden, then recurrence could be further delayed [44]. However, in order to consider additional treatments for combination therapies it is necessary to account for toxicity of the single agents, as well as toxicity resulting from their combination. The latter would impose an additional constraint in the optimization problem. Here we made no assumption regarding the toxicity of both inhibitors and therefore allowed any length of continuous targeted drug administration. Nevertheless, it is worth noting that the intermittent drug treatment we propose not only delays progression but also uses only a third of the drug, when compared to continuous treatment.

The heterogeneity our study revealed from the *in vivo* experiments highlights the importance in accounting for mouse-(human) specific micro environmental parameters to accurately capture response dynamics. This heterogeneity is often ignored in pre-clinical models, as they aim at establishing general relationships of causality between biological mechanisms. However, as our study suggests, heterogeneity can be key in explaining the variation observed across replicates of an experimental system. Furthermore, models that exploit the transient nature of EMDR must rely on individually calibrated dynamics in order to propose effective and improved treatment strategies.

Whilst this study has been focused on melanoma, our model is also applicable to the treatment of other molecularly targeted tumors, such as non-small cell lung cancer. Within the practical constraints of frequency in monitoring a patient’s systemic tumor burden and tissue characteristics, our simple model could be used to drive patient (and tumor) specific treatment strategies that target both the tumor and stroma. In addition, our approach is ideally suited to directly inform the design of adaptive therapies [45].

## Acknowledgements

The authors gratefully acknowledge useful discussions with Dr. Gary Mirams, Ross Johnston and Dr. Robert Jenkins. We also thank Dr. Mark Robertson-Tessi for critical reading and feedback on earlier versions of this manuscript.

